# Differential contribution of *Anopheles coustani* and *Anopheles arabiensis* to the transmission of *Plasmodium falciparum* and *Plasmodium vivax* in two neighboring villages of Madagascar

**DOI:** 10.1101/787432

**Authors:** Jessy Goupeyou-Youmsi, Tsiriniaina Rakotondranaivo, Nicolas Puchot, Ingrid Peterson, Romain Girod, Inès Vigan-Womas, Mamadou Ousmane Ndiath, Catherine Bourgouin

## Abstract

**Background:** Malaria is still a heavy public health concern in Madagascar. Few studies combining parasitology and entomology have been recently conducted despite the need for such information to design proper vector control measures. In a region of moderate to intense transmission of both *Plasmodium falciparum* and *Plasmodium vivax*, we conducted a combined parasitology and entomology survey in two nearby villages, across a malaria transmission season from December 2016 to April 2017.

**Methodology/Principal findings:** Community-based surveys were conducted in the two close by villages at three time points during a single malaria transmission season. *Plasmodium* carriage in the human populations was determined by Rapid Diagnostic Tests (RDTs), microscopy and real-time PCR. Anthropophilic mosquitoes were captured by human landing captures and presence of *Plasmodium* sporozoites was assessed by robust Real Time PCR. Overall human malaria prevalence was 8.0% by RDT, 4.8% by microscopy and 11.9% by PCR, mainly due to *P. falciparum* detected in 92.2% of the PCR positive samples and *Plasmodium vivax* (5.7%). No significant differences in *Plasmodium* human carriage was observed between the 2 villages at any time point. Of the 1553 anopheline mosquitoes tested, 13 were found carrying *Plasmodium* sporozoites, the majority of them being captured outdoor. The mosquito sporozoite indices were not significantly different between the two villages. However, our entomological analysis revealed that *Anopheles coustani* was the main vector in one village, being responsible of 25.5 infective bites during the whole survey, whereas it was *Anopheles arabiensis* in the other village with 15 infective bites. In addition, we found a significant higher number of endophagic *An. coustani* and *An. arabiensis* in one village compared to the other.

**Conclusions/Significance:** Despite similar human malaria prevalence in two close by villages, the entomological survey demonstrated the contribution of two different mosquito species in each village, and importantly the role of a suspected secondary malaria vector, *An. coustani*, as the main vector in one village. This, along with its higher endophagic rate in that village, highlights the importance of combining parasitology and entomology surveys for better targeting the actual local malaria vector. Such study should contribute to the malaria pre-elimination goal established under the 2018-2022 National Malaria Strategic Plan.

**Author Summary:** Malaria is still a major health concern in many countries in sub-Saharan Africa such as Madagascar. In this study, we determined the contribution of malaria vectors in the transmission of *Plasmodium* parasites in two nearby villages in an area of moderate to high malaria transmission in Madagascar. We collected, during a single malaria transmission season, parasitological data in the human population and entomological data in the mosquito population, in order to evaluate *Plasmodium* carriage in these two populations. The results showed that despite similarity in human malaria prevalence and in vector species diversity in each village, the contribution of vectors was different between the two villages. *An. arabiensis* was the major vector in Ambohitromby while it was *An. coustani* that played this role in Miarinarivo. Importantly, this study is the first that clearly demonstrates that *An. coustani* could act as a major local vector in Madagascar. Such study should help deploying adapted malaria vector control and contributing to the malaria pre-elimination goal established under the 2018-2022 National Malaria Strategic Plan.

## Introduction

Malaria remains a major health concern in Madagascar with an increase of case number in 2017 [1]. Malaria epidemiology in this country is highly heterogenous and varies according to the climatic and ecological environment that allows a stratification in bioclimatic zones and ecozones [2,3]. All four human malaria species are circulating with *Plasmodium falciparum* being the most prevalent. Across the ecozones average *P. falciparum* prevalence varies from 2 to 12 % [3] but can reach 30% in some areas [4]. Among the 26 *Anopheles* species described in the country, 6 have been reported as malaria vectors with different role according to their geographic localization and behavior [5]. Three species belong to the *Anopheles gambiae* complex: *Anopheles gambiae* sensu stricto, *Anopheles arabiensis* and *Anopheles merus*, the latter having a minor role in malaria transmission, restricted to the most southern region of Madagascar [6]. Of the two other members of the *An. gambiae* complex, *An. arabiensis* is largely prevalent in most Madagascar, and plays a major role in malaria transmission along with *Anopheles funestus* [7]. *Anopheles mascarensis*, endemic to Madagascar, and *Anopheles coustani* act as local or minor vectors [8–10].

Recent analyses surveying the evolution of malaria incidence and prevalence over 2010-2016 confirm the heavy malaria burden for the population leaving in the western part of Madagascar [3,4,11]. A particular focus has recently been brought to the Tsiroanomandidy district which constitutes a bridge area between the low transmission Central Highlands and the high endemic western region where both *P. falciparum* and *P. vivax* circulate. In this area, both malaria prevalence and *Anopheles* species distribution have been documented [7,12,13]. Such information is critical for adapting the best relevant strategies for interrupting malaria transmission toward malaria elimination which has been set up on the agenda of the 2018-2022 Malagasy Malaria Strategic Plan as geographically progressive elimination. Contributing to this goal, here we report a study conducted in the Maevatanana district which is located in the northwestern ecozone of Madagascar that faces high malaria burden due to both *P. falciparum* and *P. vivax*. To our knowledge, no combined parasitological and entomological survey has ever been reported in that region. The study was specifically conducted in Andriba, a rural commune located at the transition between the western fringe of the Central Highlands (low malaria prevalence) and the north western ecozone (moderate to high prevalence) according to Howes *et al.* [3]. Survey of malaria prevalence in the population of two villages 1.5 km apart revealed a high level of submicroscopic carriage of *Plasmodium*, similar to the one reported in the Tsiroanomandidy district [13]. Importantly, this study showed that in these close by villages *Plasmodium* transmission was due to two different mosquito species, *An. arabiensis* and *An. coustani*. The role of *An. coustani* as the main malaria vector in one village was quite surprising as this mosquito species is being known as a zoophilic and exophilic species in most places of Madagascar where it has been found. Our data revealed a high proportion of indoor biting and anthropophilic *An. coustani* mosquitoes. These results highlight the needs to better investigate the role of each malaria vector in every malaria transmission hotspot with appropriate tools for developing adapted vector control measure towards malaria elimination.

## Methods

### Study Design and Setting

The study was conducted in two villages of the rural commune of Andriba (Maevatanana district, Madagascar) which is located in the tropical northwest region of Madagascar. Andriba is characterized by a dry season that generally lasts from April to October and a rainy season from November to April; the average annual temperature is 24°C and the average annual rainfall is 1828 mm [14]. The two villages, Ambohitromby (17°34’23.7”S, 46°55’21.4”E) and Miarinarivo (17°33’56.7”S, 46°55’10.8”E) are located 1.5 kilometers apart and 6 kilometers from Andriba town hall (Fig 1).

**Fig 1.**
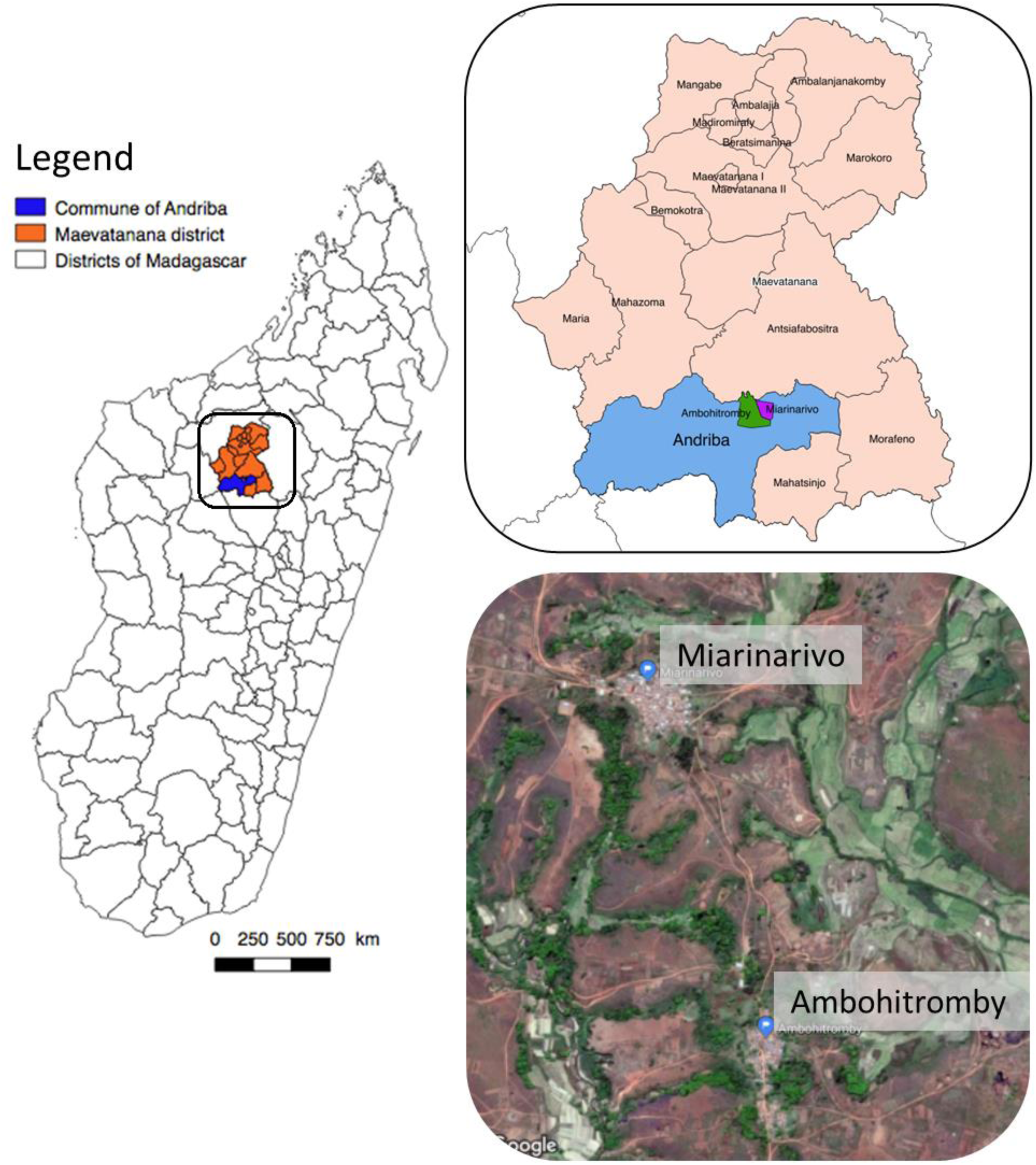
Study site. The map of Madagascar is depicted on the left panel with a focus on the Andriba region presented in more details in the upper right panel. The bottom right panel is a Google view of the studied villages, Ambohitromby and Miarinarivo, located 1.5 km apart.

Parasitological and entomological data were collected at 3 time points during a single malaria transmission season: at the onset of the season (December 2016, labelled T1), mid-season (February 2017, labelled T2) and late-season (April 2017, labelled T3). The latter time point was estimated to correspond with the cessation of malaria transmission, but this depends upon annual climatic variation. Additional parasitological data were recorded in March 2016 and in March 2018 as part of active malaria parasite surveillance in school children (Bourgouin and coll, unpublished).

The study was approved by the Malagasy Ethical Committee of the Ministry of Health (agreements N°122-MSANP/CE–2015 and N°141-MSANP/CE–2014). Prior to carrying out study procedures, individual written informed consent was obtained from all study participants, or their parents or legal guardians in the case of minors.

### Malaria prevalence in the Human population

#### Blood sample collection

Blood samples were collected from children and adults without malaria symptoms. Study participants included village residents among which volunteers involved in Human landing catches (HLCs) and their family members. Blood samples were obtained by finger prick to perform RDTs (SD BIOLINE Malaria Ag P.f/Pan), thick and thin blood smears, and blood spots on filter papers (standard Whatman 3MM filter paper). The Bioline RDT enabled specific detection of *P. falciparum*, and any of *P. vivax, P. ovale* and *P. malariae*, as all four species are present in Madagascar. *Plasmodium* species, parasite stages and quantification were further determined by microscopic observation of the blood smears stained with 10% Giemsa, using a light microscope (100X). A thin blood smear slide was declared malaria-negative when *Plasmodium* parasites were not detected after examination of 100 high power microscopic fields. Slides were read for asexual parasites and gametocytes, enumerated against 500 leucocytes and expressed as density/μL assuming an average leucocyte count of 8,000/μL of blood (data not shown). Individuals with a positive RDT were treated with Artemisinin-based combination therapy (ACT), according to national guidelines.

#### Submicroscopic detection of *Plasmodium* parasites by PCR using Dried Blood Spots

*Plasmodium* carriage prevalence was also assessed by PCR which is more sensitive than both RDT and microscopy for parasite detection [15]. Dried blood spots were lysed overnight at 4°C with 150 µl per well of 1X HBS buffer (Hepes Buffer Saline) supplemented with 0.5% Saponin, final concentration. Samples were then washed twice with 1X PBS and DNA was extracted with Instagene® Matrix resin (Bio-Rad, France) according to manufacturer’s instructions. Molecular detection and species identification of *Plasmodium* parasites were performed in 2 steps as previously described by Canier *et al.* [16]. *Plasmodium* parasites were first detected by a real-time PCR screening with genus-specific primers targeting the *Plasmodium* cytochrome b gene. Then, *Plasmodium* species identification was performed on DNA samples identified as positive for *Plasmodium* using a nested real-time PCR assay.

### Entomological data

#### Mosquito collection

Anthropophilic female mosquitoes were collected by HLCs and indoor Pyrethrum spray catches (PSCs), following WHO protocols [17]. At each survey time point during the malaria season, adult volunteers performed HLCs from 18h:00 to 06h:00 in 2 houses per night, on 3 consecutive nights. One volunteer was sitting inside and another outside and they changed places every hour. Two volunteers worked between 18h:00 to 24h:00, and were replaced by 2 others who worked from 24h:00 to 06h:00. Therefore, for each house, the 4 volunteers conducting HLCs represented 2 human-nights (HNs); in total we collected HLCs data from 12 HNs at each time point, in each village. The whole study corresponds to 72 HNs. PSCs were conducted in 5 houses/day/village, choosing houses that were not used for HLCs and in which no insecticide or repellent has been used during the previous week. PSCs were done on 3 consecutive days, on each morning (from 6 to 8 am) following the HLCs. No insecticide residual spraying had occurred in the villages from at least 2016 till 2018. PSCs were performed using a pyrethroid mixture of Prallethrin, Tetramethrin, and Deltamethrin. Spraying was performed from outside of the houses, into openings, holes in walls and eaves, then in the rooms following WHO procedures. Knockdown mosquitoes were then collected by hand picking. The houses were of typical Malagasy construction common in the rural areas of Maevatanana district: thatched roofs, adobe walls and composed of 1 to 2 rooms (Fig 2). For both HLCs and PSCs, houses were chosen randomly in the villages and with no repetition.

**Fig 2.**
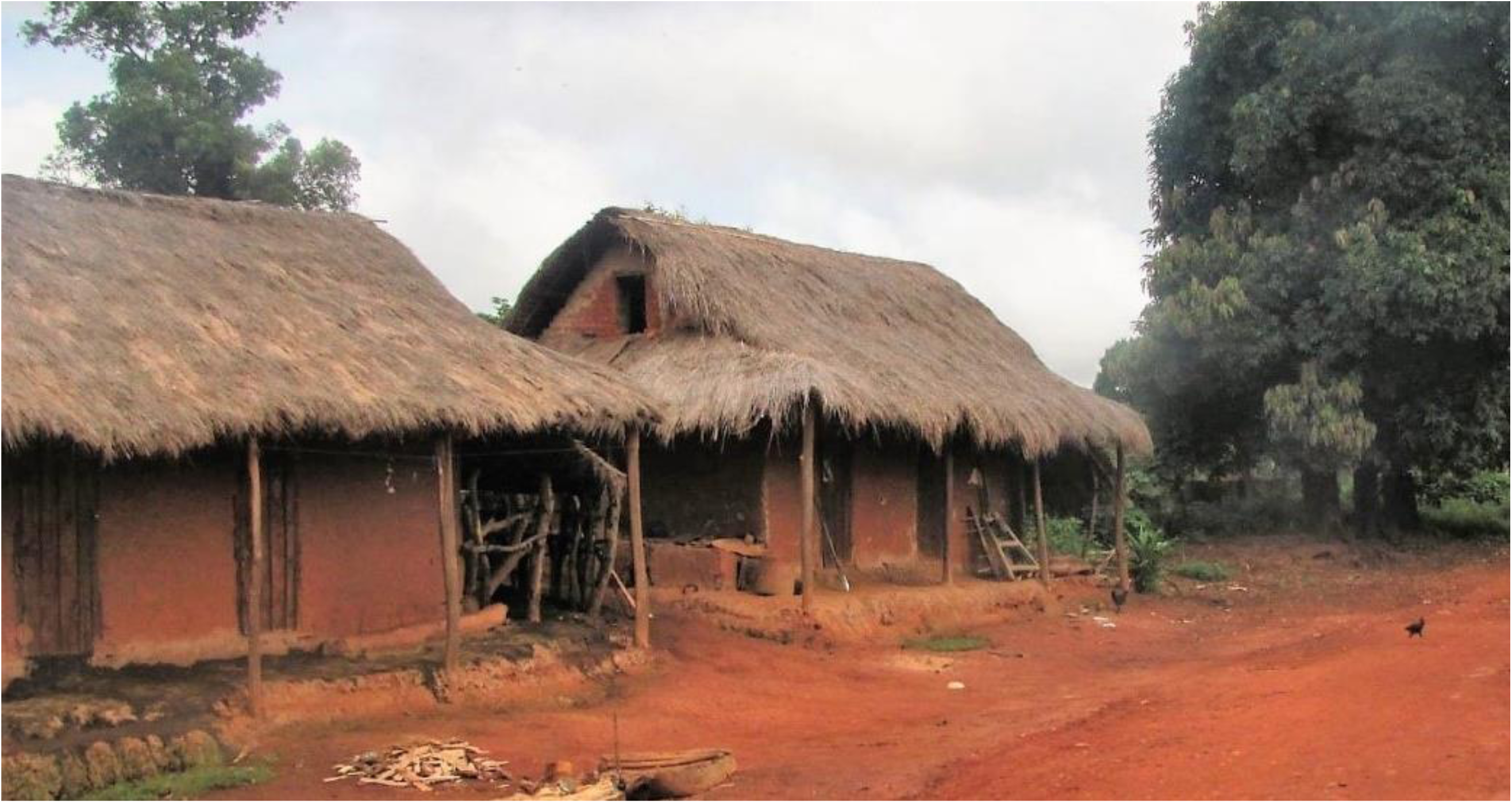
Typical Malagasy houses in Andriba rural area. The houses are built with adobe walls and thatched roofs, and usually composed of one or two rooms. The picture was taken in the village of Ambohitromby located in the rural commune of Andriba, Madagascar.

#### Mosquito identification

All mosquitoes were identified morphologically using the determination keys of Grjebine[18] and De Meillon [19]. To discriminate *An. gambiae* from *An. arabiensis*, a TaqMan assay was used targeting the intergenic spacer region of rDNA as described by Walker *et al*. [20], following the initial work of Scott *et al* [21]. Primers and sequences used are listed in S1 Table. PCR reactions (20µL) contained 5µl of genomic DNA (see extraction procedure below), 4µL of 5x HOT FIREPol® Probe qPCR Mix Plus /no ROX (Solis Biodyne), 300nM of each primer and 200nM of each probe. Reactions were run on a StepOnePlus (Applied Biosystems) using the following temperatures: an initial step at 95°C for 10min, followed by 40 cycles of denaturation at 95°C for 20sec and annealing at 60°C for 1min.

### *Plasmodium* detection in *Anopheles* mosquitoes

#### DNA extraction and Quality control

Genomic DNA from *Anopheles* head-thorax was extracted using the DNAzol® Reagent (ThermoFisher Scientific). Briefly, the head-thorax from each mosquito was put individually in a tube; care was taken to rinse the dissecting equipment in ethanol 70% between each mosquito. A volume of 150µL of DNazol was added and mosquito tissues were crushed using individual conical plastic pestle. DNA was then extracted following the manufacturer’s protocol. After precipitation, the DNA pellet was suspended in a final volume of 50µL of nuclease-free water. DNA quality was controlled using a SYBR Green real-time PCR assay targeting the ribosomal S7 protein encoding gene. This gene is highly conserved among species belonging to the same genus. Primers previously designed against the *An. gambiae* gene [22] were aligned against all available *Anopheles* S7 sequences to ensure that those primers will efficiently amplified the S7 gene fragment from any *Anopheles* captured in the field. Amplification conditions were validated on a subset of laboratory and field mosquito samples including *Anopheles, Cellia* and *Nyssorhynchus* subgenera (not shown). Amplification using the PowerSYBER® Green Master mix (Applied biosystems) was performed as follows: an initial step at 95°C for 15min, followed by 40 cycles of denaturation at 95°C/45 sec, annealing at 55°C/30sec and elongation at 60°C/45sec. Specificity of the amplification was assessed by viewing the melting curves.

#### *Plasmodium* detection

The detection of human *Plasmodium* gDNA in mosquitoes was performed in 2 steps. The first step used a TaqMan PCR assay targeting a region of the 18S rRNA gene conserved among the human infecting *Plasmodium* species. For this assay, we used primers and probe previously described [23] using a MGB probe as in Taylor *et al.*[24]; this combination was previously validated for *Plasmodium* detection in mosquitoes [25]. The *Plasmodium* TaqMan probe was labelled with 5’NED. PCR reactions (20µL) contained 5µL of mosquito genomic DNA, 4µL of 5x HOT FIREPol® Probe qPCR Mix Plus/no ROX (Solis Biodyne), 300nM of each primer and 200nM of probe. Reactions were run on a StepOnePlus (Applied Biosystems) using the following conditions: an initial step at 95°C/10min, followed by 50 cycles of denaturation at 95°C/20sec and annealing and elongation at 60°C/1min. *P. falciparum* genomic DNA extracted from NF54 parasite cultures was used as positive control. Any sample with amplification signal before the 38th cycle was considered positive. For high through put screening, a pool strategy was used [26]; equal volumes of genomic DNA (extracted as described above) from 6 mosquitoes of the same species were pooled. The *Plasmodium* TaqMan assay was run using 5µL of each DNA pool in triplicate. Mosquitoes from positive pools were then analyzed individually as for pools. All positive samples in the TaqMan assay were then analyzed for the identification of *P. falciparum* and *P. vivax* species. For each positive sample, 2 distinct real-time SYBR Green PCR assays were done using species-specific primers targeting the cytochrome b gene. Purified gDNA from *P. falciparum* and *P. vivax* were used as positive control. Each reaction was run in triplicate. The real-time PCR conditions were as previously described [16]: an initial step at 95°C for 15min, followed by 40 cycles of denaturation at 95°C/20sec and annealing + elongation at 60°C/1min. Sequences of the primers and TaqMan probes used for the *Plasmodium* detection in *Anopheles* mosquitoes are listed in S1 Table.

#### Statistical analysis

Stata 15 (StataCorp. 2017. Stata Statistical Software: Release 15. College Station, TX: StataCorp LLC) were used for statistical analysis. Pearson’s Chi-squared test was used to compare malaria prevalence and the proportion of collected anopheline species between the two villages. The Fisher’s exact was used when sample size was small (< 30).. *P* values < 0.05 were considered to be statistically significant.

## Results

### *Plasmodium* carriage in the Human population

The parasitological survey involved 380 individuals ranging from 5 months to 68 years old (S2 Table). A total of 590 samples (351 in Ambohitromby and 239 in Miarinarivo) were analyzed by RDT, microscopy and real-time PCR. Human malaria prevalence was 8.0% by RDT, 4.8% by microscopy and 11.9% by real-time PCR over the whole study (Table 1). Whatever the diagnosis method, the prevalence of *Plasmodium* infections was similar in the two villages at each time point with a peak at T2 for both RDT and PCR. Except for T3 in Miarinarivo, the PCR technique, as expected, was able to detect a greater number of parasite carriers over RDT and microscopy, revealing a substantial proportion of submicroscopic parasite carriers. The lower proportion of PCR positive samples at T3 Miarinarivo might result from inadequate conservation of the blood spots before PCR processing.

**Table 1.** Prevalence of *Plasmodium* infections in asymptomatic individuals obtained from RDT, Microscopy and real-time PCR.

|  |  | RDT |  |  |  | Microscopy |  |  |  | Real-time PCR |  |  |  |
| --- | --- | --- | --- | --- | --- | --- | --- | --- | --- | --- | --- | --- | --- |
|  |  | T1 | T2 | T3 | Total | T1 | T2 | T3 | Total | T1 | T2 | T3 | Total |
| **Ambohitromby** | Sample size | 173 | 79 | 99 | **351** | 173 | 79 | 99 | **351** | 173 | 79 | 99 | **351** |
|  | Positive | 13 | 10 | 5 | **28** | 9 | 3 | 3 | **15** | 20 | 12 | 10 | **42** |
|  | Prevalence | 7.5% | 12.7% | 5.1% | **8.0%** | 5.2% | 3.8% | 3.0% | **4.3%** | 11.6% | 15.2% | 10.1% | **12.0%** |
| **Miarinarivo** | Sample size | 122 | 65 | 52 | **239** | 122 | 65 | 49 | **236** | 122 | 65 | 52 | **239** |
|  | Positive | 9 | 7 | 3 | **19** | 7 | 3 | 3 | **13** | 14 | 12 | 2 | **28** |
|  | Prevalence | 7.4% | 10.8% | 5.8% | **7.9%** | 5.7% | 4.6% | 6.1% | **5.5%** | 11.5% | 18.5% | 3.8% | **11.7%** |
| Prevalence of *Plasmodium* infections in the two villages |  | 7.5% | 11.8% | 5.3% | **8.0%** | 5.4% | 4.2% | 4.1% | **4.8%** | 11.5% | 16.7% | 7.9% | **11.9%** |
| *P* values (Chi2 test) between the two villages |  | 0.965 | 0.727 | 0.851 | **0.990** | 0.842 | 0.807 | 0.369 | **0.491** | 0.982 | 0.600 | 0.177 | **0.926** |

The same individuals were involved during the transversal parasitological study including the three methods of malaria diagnostic. In Ambohitromby, a total of 173, 79 and 99 samples were analyzed for T1, T2 and T3 respectively. In Miarinarivo, a total of 122, 65 and 52 samples were analyzed for T1, T2 and T3, respectively, except at T3 were only 49 smears were read as 3 slides were unreadable.

Among the 70 positive samples assessed by real-time PCR, 84.3% were due to *P. falciparum*, 5.7% to *P. vivax*, 1.4% to *P. malariae* and 8.6% to mixed infections always involving *P. falciparum* (Table 2). All mixed infections were observed in Ambohitromby.

**Table 2.**
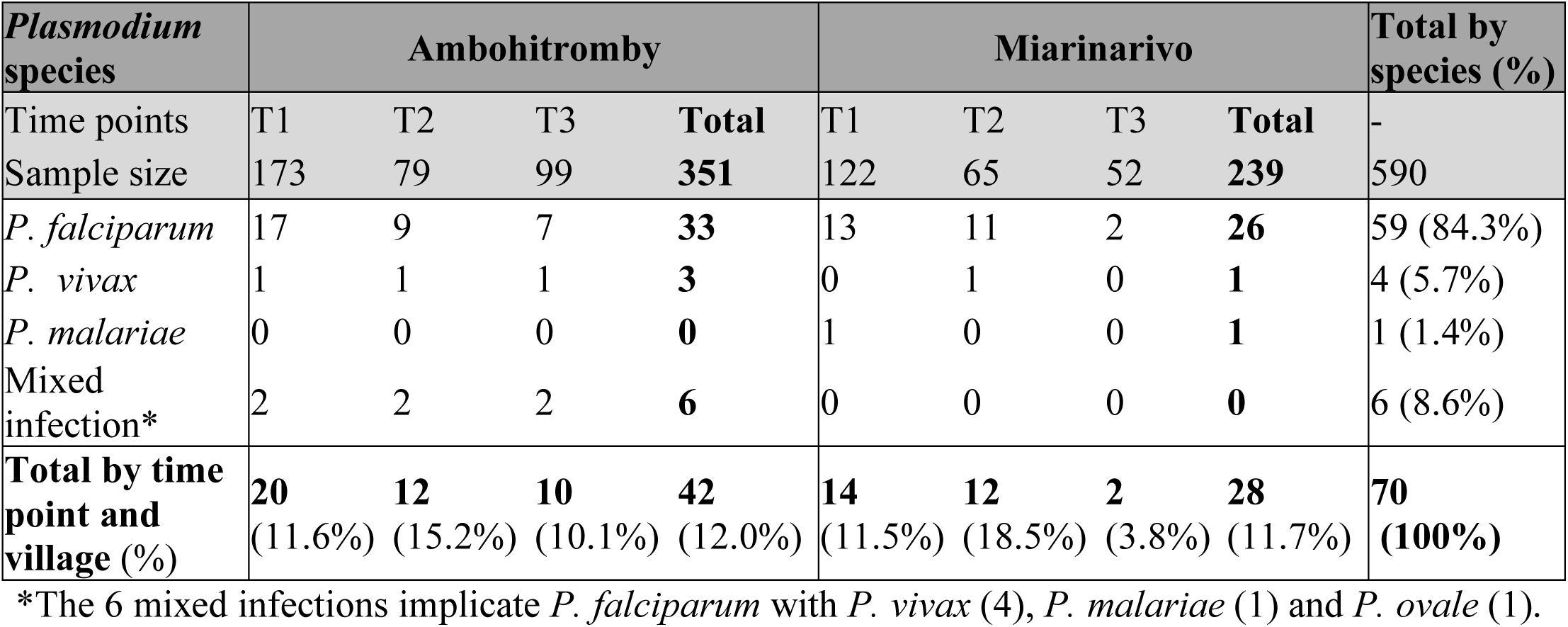
*Plasmodium* species detected by real-time PCR in asymptomatic individuals.

| ***Plasmodium* species** | **Ambohitromby** |  |  |  | **Miarinarivo** |  |  |  | **Total by species (%)** |
| --- | --- | --- | --- | --- | --- | --- | --- | --- | --- |
| Time points | T1 | T2 | T3 | **Total** | T1 | T2 | T3 | **Total** | - |
| Sample size | 173 | 79 | 99 | **351** | 122 | 65 | 52 | **239** | 590 |
| *P. falciparum* | 17 | 9 | 7 | **33** | 13 | 11 | 2 | **26** | 59 (84.3%) |
| *P. vivax* | 1 | 1 | 1 | **3** | 0 | 1 | 0 | **1** | 4 (5.7%) |
| *P. malariae* | 0 | 0 | 0 | **0** | 1 | 0 | 0 | **1** | 1 (1.4%) |
| Mixed infection* | 2 | 2 | 2 | **6** | 0 | 0 | 0 | **0** | 6 (8.6%) |
| **Total by time point and village (%)** | **20**<br>(11.6%) | **12**<br>(15.2%) | **10**<br>(10.1%) | **42**<br>(12.0%) | **14**<br>(11.5%) | **12**<br>(18.5%) | **2**<br>(3.8%) | **28**<br>(11.7%) | **70**<br>**(100%)** |

\*The 6 mixed infections implicate *P. falciparum* with *P. vivax* (4), *P. malariae* (1) and *P. ovale* (1).

### Vector species and Behavior

#### Mosquito Abundance and Aggressiveness

In total, 2407 mosquitoes were collected during 72 HNs in Ambohitromby and Miarinarivo. As presented in Table 3, *Anopheles* was the most abundant mosquito genus collected (68.55%, n=1650) followed by *Culex* (26.87%, n=647), *Mansonia* (3.49%, n=84), *Aedes* (0.99%, n=24) and *Coquillettidia* (0.08%, n=2). Among *Anopheles, An. coustani* was by far the most abundant representing 52.99% (751/1417) of the known potential malaria vectors in Madagascar, followed by *An. arabiensis* (28.93%, 410/1417), *An. funestus* (12.84%, 182/1417) and *An. mascarensis* (4.66%, 66/1417); *An. gambiae* was barely represented (0.56%, 8/1417). Detailed data covering all collected species are presented in S3 Table.

**Table 3.** Mosquitoes collected by HLCs in Ambohitromby and Miarinarivo over the 3 survey time periods.

| Mosquito species | Ambohitromby |  |  |  | Miarinarivo |  |  |  | Grand Total |
| --- | --- | --- | --- | --- | --- | --- | --- | --- | --- |
|  | T1 | T2 | T3 | Total | T1 | T2 | T3 | Total |  |
| *Anopheles arabiensis*<sup>a,b</sup> | 64 | 207 | 10 | 281 | 16 | 81 | 32 | 129 | 410 |
| *Anopheles gambiae*<sup>a,b</sup> | 0 | 4 | 0 | 4 | 1 | 2 | 1 | 4 | 8 |
| *Anopheles funestus*<sup>a</sup> | 34 | 26 | 40 | 100 | 9 | 14 | 59 | 82 | 182 |
| *Anopheles mascarensis*<sup>a</sup> | 24 | 13 | 12 | 49 | 5 | 2 | 10 | 17 | 66 |
| *Anopheles coustani*<sup>a</sup> | 75 | 42 | 162 | 279 | 66 | 97 | 309 | 472 | 751 |
| *Anopheles squamosus/cydippis* | 13 | 43 | 29 | 85 | 8 | 40 | 15 | 63 | 148 |
| *Anopheles rufipes* | 9 | 10 | 0 | 19 | 5 | 11 | 9 | 25 | 44 |
| *Anopheles maculipalpis* | 7 | 9 | 0 | 16 | 10 | 9 | 5 | 24 | 40 |
| *Anopheles pretoriensis* | 0 | 0 | 0 | 0 | 0 | 0 | 1 | 1 | 1 |
| **Total Anopheles** | **226** | **354** | **253** | **833** | **120** | **256** | **441** | **817** | **1650** |
| *Culex antennatus* | 30 | 258 | 4 | 292 | 49 | 99 | 25 | 173 | 465 |
| *Culex quinquefasciatus* | 5 | 103 | 0 | 108 | 15 | 23 | 19 | 57 | 165 |
| *Culex bitaeniorhynchus* | 0 | 0 | 0 | 0 | 1 | 1 | 0 | 2 | 2 |
| *Culex univittatus* | 4 | 0 | 0 | 4 | 0 | 0 | 0 | 0 | 4 |
| *Culex decens* | 1 | 0 | 0 | 1 | 2 | 0 | 0 | 2 | 3 |
| *Culex giganteus* | 0 | 0 | 0 | 0 | 2 | 4 | 2 | 8 | 8 |
| **Total Culex** | **40** | **361** | **4** | **405** | **69** | **127** | **46** | **242** | **647** |
| *Mansonia uniformis* | 8 | 9 | 3 | 20 | 5 | 25 | 34 | 64 | 84 |
| **Total Mansonia** | **8** | **9** | **3** | **20** | **5** | **25** | **34** | **64** | **84** |
| *Aedes skusea* | 5 | 0 | 0 | 5 | 3 | 0 | 0 | 3 | 8 |
| *Aedes vittatus* | 1 | 1 | 0 | 2 | 0 | 0 | 0 | 0 | 2 |
| *Aedes tiptoni* | 1 | 0 | 0 | 1 | 8 | 0 | 1 | 9 | 10 |
| *Aedes albopictus* | 0 | 1 | 0 | 1 | 1 | 1 | 0 | 2 | 3 |
| *Aedes circumlateolus* | 0 | 0 | 0 | 0 | 0 | 0 | 1 | 1 | 1 |
| **Total Aedes** | **7** | **2** | **0** | **9** | **12** | **1** | **2** | **15** | **24** |
| *Coquilleltidia grandidieri* | 0 | 0 | 0 | 0 | 0 | 0 | 2 | 2 | 2 |
| **Total Coquilleltidia** | **0** | **0** | **0** | **0** | **0** | **0** | **2** | **2** | **2** |
| **Grand Total** | **281** | **726** | **260** | **1267** | **206** | **409** | **525** | **1140** | **2407** |

<sup>a</sup> known potential malaria vectors in Madagascar. <sup>b</sup> *An. arabiensis* and *An. gambiae* were identified by TaqMan assay among all *An. gambiae* s.l. collected (see M&M).

The aggressive density *ma* of the known potential malaria vectors, (except *An. gambiae*, n=8) over time and for each village is depicted in Figure 3. Overall *Anopheles* aggressiveness varies in each village over the time course of the survey, from 8 bites per man and per night (b/m/n) to 34.3 b/m/n. However, the bite frequency was higher at T2 (middle of the malaria transmission season) in Ambohitromby, while being the highest at T3 (vanishing of the malaria transmission season) in Miarinarivo. Strinkingly, *An. arabiensis* was the most aggressive species in Ambohitromby with 17.3 b/m/n at T2 while it was *An. coustani* with 25.9 b/m/n in Miarinarivo at T3. Nevertheless, *An. coustani* exhibited the highest biting frequency in both villages at T3, which corresponds to the time where the rice reached full maturity providing high shade on the padding fields suitable for *An. coustani* larval development.

**Fig 3.**
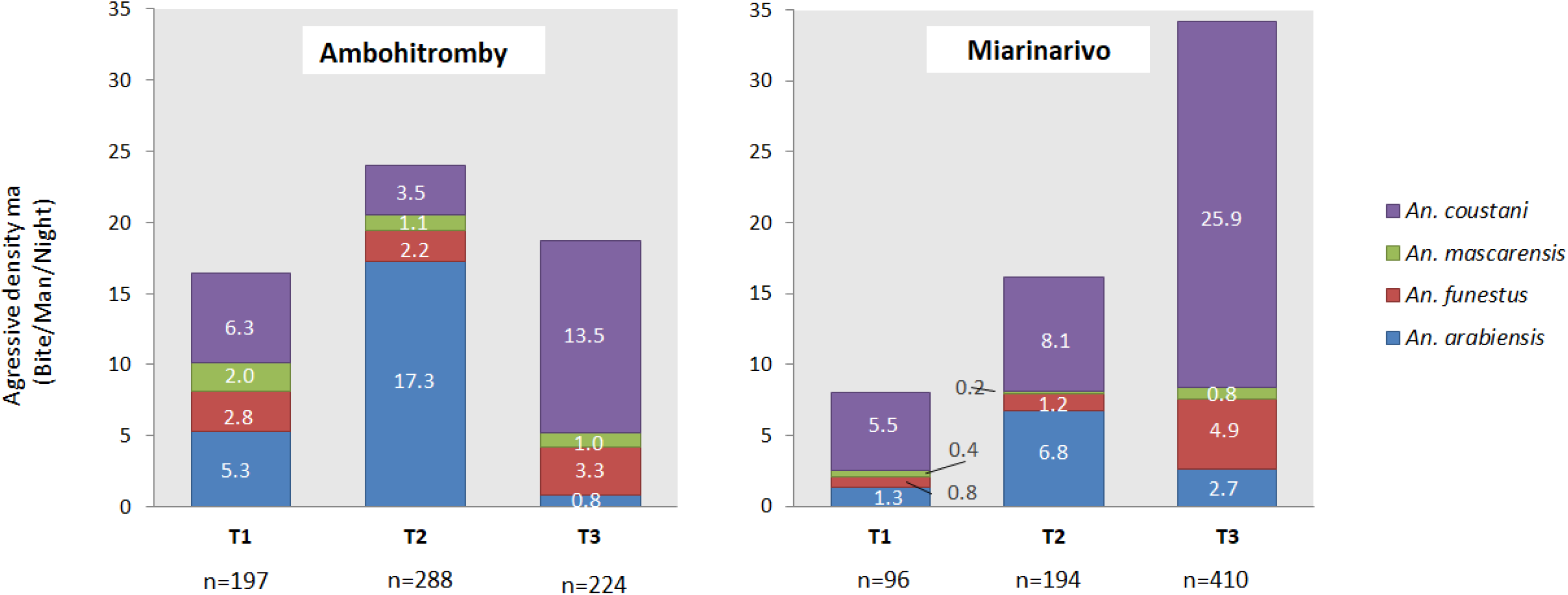
Aggressive density of the malaria vectors in Ambohitromby and Miarinarivo at the three time points. n= number of mosquitoes collected by HLCs, both indoor and outdoor. Light numbers within the graphs indicate the mean bite per man and per night for each of the four *Anopheles* species.

Looking at the hourly aggressiveness of the four potential malaria vectors across time-points and the two villages, the 4 mosquito species globally bite all night long (Fig 4). There are however variations in the biting pattern across the night. In Miarinarivo, the *An. coustani* population collected at T1 and T3 tend to bite early at night compared to the population collected at T2 and the one collected at T3 in Ambohitromby. Of note is the bi-modal hourly biting rate observed at T2 in Miarinarivo for both *An. arabiensis* and *An. coustani* and for which no rational explanation can be proposed beside a human bias during the HLCs or occurrence of strong winds.

**Fig 4.**
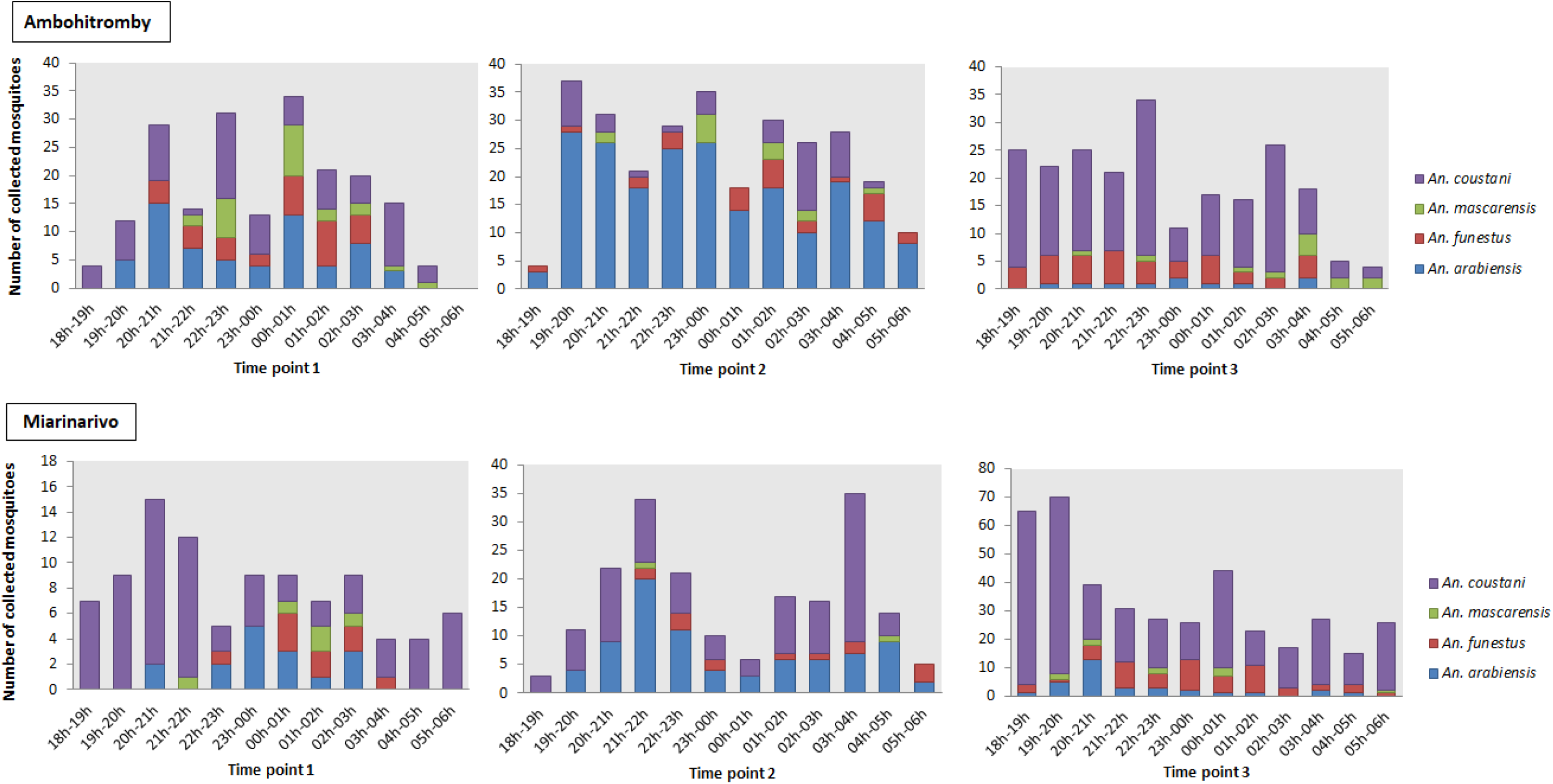
Hourly aggressiveness of malaria vectors in Ambohitromby and Miarinarivo at the three time points. Data represent both indoor and outdoor HLCs collected mosquitoes. Note that the scales in Miarinarivo at time points 1 and 3 are different from the others.

#### Endophagy and Endophily rates

Endophagy rates, representing the proportion of mosquitoes collected indoor over the total number of mosquitoes collected indoor and outdoor by HLCs, are summarized in Table 4 for the four more abundant potential malaria vector species, according to the time-course of the survey and each village. Among those *Anopheles* species, *An. funestus* exhibited the highest mean endophagic rate in both villages, not departing from its known endophagic behavior [27–30]. Of note is its high endophagic rate (77.50% and 88.89%) at T3 in Ambohitromby and T1 in Miarinarivo, respectively. Such high endophagic rates have been reported in other places in Madagascar [7]. Conversely, *An. mascarensis* exhibited the lowest endophagic rates with mean values of 8.12% ± 6.81% and 10.00 ± 8.16% in Ambohitromby and Miarinarivo, respectively. Those values are in accordance with previous reports [7] collated in Goupeyou-Youmsi *et al.* (in preparation). By contrast, *An. arabiensis* and *An. coustani* showed significant different endophagic rates whether collected in Ambohitromby or Mirarinarivo. Indeed, both species, known as zoo-anthropophilic species, exhibited a higher significant endophagic rate in Miarinarivo compared to Ambohitromby. The mean endopaghagic rate for *An. arabiensis* was 27.79 ± 2.91% in Ambohitromby versus 47.76 ± 8.00% in Miarinarivo (significant difference P<0.05, Pearson Chi2), and 6.74 ± 6.55% in Ambohitromby versus 31.95 ± 5.00% in Miarinarivo (significant difference P<0.05, Pearson Chi2) for *An. coustani.* This observation might suggest the presence of different populations for both species in each village.

**Table 4.** Proportion of the malaria vectors collected biting indoor by HLCs (Endophagic rate).

| Species | Ambohitromby |  |  |  | Miarinarivo |  |  |  |
| --- | --- | --- | --- | --- | --- | --- | --- | --- |
| Time points | T1 | T2 | T3 | Mean<br>± SD | T1 | T2 | T3 | Mean<br>± SD |
| *Anopheles arabiensis* | 29.69%<br>(19/64) | 23.67%<br>(49/207) | 30.00%<br>(3/10) | **27.79**<br>± **2.91%** | 56.25%<br>(9/16) | 37.04%<br>(30/81) | 50.00%<br>(16/32) | **47.76**<br>± **8.00%** |
| *Anopheles funestus* | 41.18%<br>(14/34) | 50.00%<br>(13/26) | 77.50%<br>(31/40) | **56.23**<br>± **15.47%** | 88.89%<br>(8/9) | 50.00%<br>(7/14) | 49.15%<br>(29/59) | **62.68**<br>± **18.54%** |
| *Anopheles mascarensis* | 16.67%<br>(4/24) | 7.69%<br>(1/13) | 0.00%<br>(0/12) | **8.12**<br>± **6.81%** | 20.00%<br>(1/5) | 0.00%<br>(0/2) | 10.00%<br>(1/10) | **10.00**<br>± **8.16%** |
| *Anopheles coustani* | 16.00%<br>(12/75) | 2.38%<br>(1/42) | 1.85%<br>(3/162) | **6.74**<br>± **6.55%** | 31.82%<br>(21/66) | 38.14%<br>(37/97) | 25.89%<br>(80/309) | **31.95**<br>± **5.00%** |

SD: Standard Deviation. Numbers in parenthesis represent the number of mosquitoes collected indoor over the total number of mosquitoes collected indoor and outdoor. *An. gambiae* was not taken into account in this table due to its low number.

From 17 PSCs, a total of 70 mosquitoes were collected resting indoor (S4 Table). Among those 42.10% were *An. funestus*, 31.57% *An. arabiensis*, 15.78% *An. coustani* and 10.52% *An. mascarensis* in Ambohitromby, while only *An. funestus* (96.15%) and *An. mascarensis* (3.84%) were collected resting indoor in Miarinarivo. The absence of indoor resting *An. arabiensis* and *An. coustani* in Miarinarivo despite their abundance, might also advocate for the presence of different populations for both species in each village.

#### *Plasmodium* carriage in anopheline mosquitoes and EIR

Among the 1715 anopheline mosquitoes captured by HLCs (n=1650) and PSCs (n=65), 1553 were tested for presence *Plasmodium* sporozoites by TaqMan assay and Real Time PCR. As described in M&M, DNA was first extracted from head-thorax of individual mosquitoes and its quality assessed by amplification of the S7 gene. Using an equal volume of gDNA from at most 6 mosquitoes of the same species, 261 pools were assembled and tested for the presence of *Plasmodium* DNA, using the *Plasmodium* Taqman assay. Twenty-three (23) pools were found containing *Plasmodium* DNA. Deconvolution of each positive pool to individual mosquito revealed that 28 mosquitoes had *Plasmodium* DNA in their head-thorax, corresponding to mosquitoes captured by HLCs only. However, the SYBR Green assay for *P. falciparum*/*P. vivax* species detection was conclusive only for 13 out 28 mosquitoes carrying either *P. falciparum* or *P. vivax* parasite (Table 5). The 15 remaining TaqMan *Plasmodium-* positive mosquitoes were either infected by *P. malariae* that is circulating in the villages, or the chosen TaqMan *Plasmodium* assay cut-off was not stringent enough. Additionally, it cannot be excluded that non-human malaria parasites, such as Lemur malaria parasites were detected in those mosquitoes as the TaqMan primer-probe combination used was conserved among *Plasmodium sp.* 18S gene. Overall, 9 mosquitoes were positive for *P. falciparum* and 4 for *P. vivax*, and belong to 3 anopheline species: *An. funestus An. arabiensis* and *An. coustani*, with the latter species being the more frequently infected one (Table 5). The sporozoite index (SI) varies from 0 to 1.4% according to the *Anopheles* species, giving an overall SI of 0.84% when taking into account all *Anopheles* species tested, including mosquitoes captured by HLCs and PSCs. Of note, *An. rufipes*, an anopheline species which is not known being a malaria vector in Madagascar, was found positive by the TaqMan *Plasmodium* assay (2/40). As there is increased reports on its role in malaria transmission in other countries [31–33], it might be worth to include this species for *Plasmodium* sporozoite carriage in future surveillance programs.

**Table 5.** *Plasmodium* carriage in anopheline mosquitoes analyzed in pool and individually.

| Species | Total screened | Pools analyzed | Positive pools | Positive mosq. | Positive in species screening | *Plasmodium* species (number) | Sporozoitic Index (SI) |
| --- | --- | --- | --- | --- | --- | --- | --- |
| *An. arabiensis* | 374 | 60 | 8 | 9 | 3 | *Pf* (1), *Pv* (2) | 0.80% |
| *An. funestus* | 213 | 36 | 3 | 3 | 3 | *Pf* (2), *Pv* (1) | 1.40% |
| *An. coustani* | 715 | 122 | 10 | 14 | 7 | *Pf* (6), *Pv* (1) | 0.98% |
| *An. rufipes* | 40 | 7 | 1 | 2 | 0 | - | 0% |
| *An. mascarensis* | 59 | 10 | 1 | 0 | 0 | - | 0% |
| *An. squamosus* | 117 | 20 | 0 | 0 | 0 | - | 0% |
| *An. maculipalpis* | 35 | 6 | 0 | 0 | 0 | - | 0% |
| **Total** | **1553** | **261** | **23** | **28** | **13** | ***Pf* (9), *Pv* (4)** | **0.84%** |

mosq. mosquitoes; *Pf*: *P. falciparum*; *Pv*: *P. vivax*. Mosquitoes from PSCs are included among the 1553.

When looking at the Entomological Inoculation Rate (EIR) a proxy for malaria transmission, it appears that *An. arabiensis* and *An. coustani* are the vector species that contribute most to malaria transmission, but with striking differences between the two villages and over time (Table 6). Indeed, in Ambohitromby *An. arabiensis* was the main vector at the beginning of transmission season (T1) with an EIR of 0.18 ib/m/n, followed by *An. coustani* at T2 (0.14 ib/m/n) whereas in Miarinarivo *An. coustani* was the main vector at both T2 and T3 (EIR of 0.26 ib/m/n each) with no role of *An. arabiensis* in that village. Plotting the relative abundance of *An. arabiensis* and *An. coustani*, and their respective EIR over time clearly shows that despite similar abundance in Ambohitromby and Miarinarivo at T3, *An. coustani* acted as a malaria vector only in Miarinarivo (Fig. 5). Conversely, in that village despite similar abundance at T1 and T3, *An. coustani* contributed to malaria transmission at T3 only. This might be a consequence of a specificity of the *An. coustani* population as hypothesized from the observed higher endophagic behavior of this mosquito species in Miarinarivo compared to Ambohitromby (see Table 4).

**Table 6.**
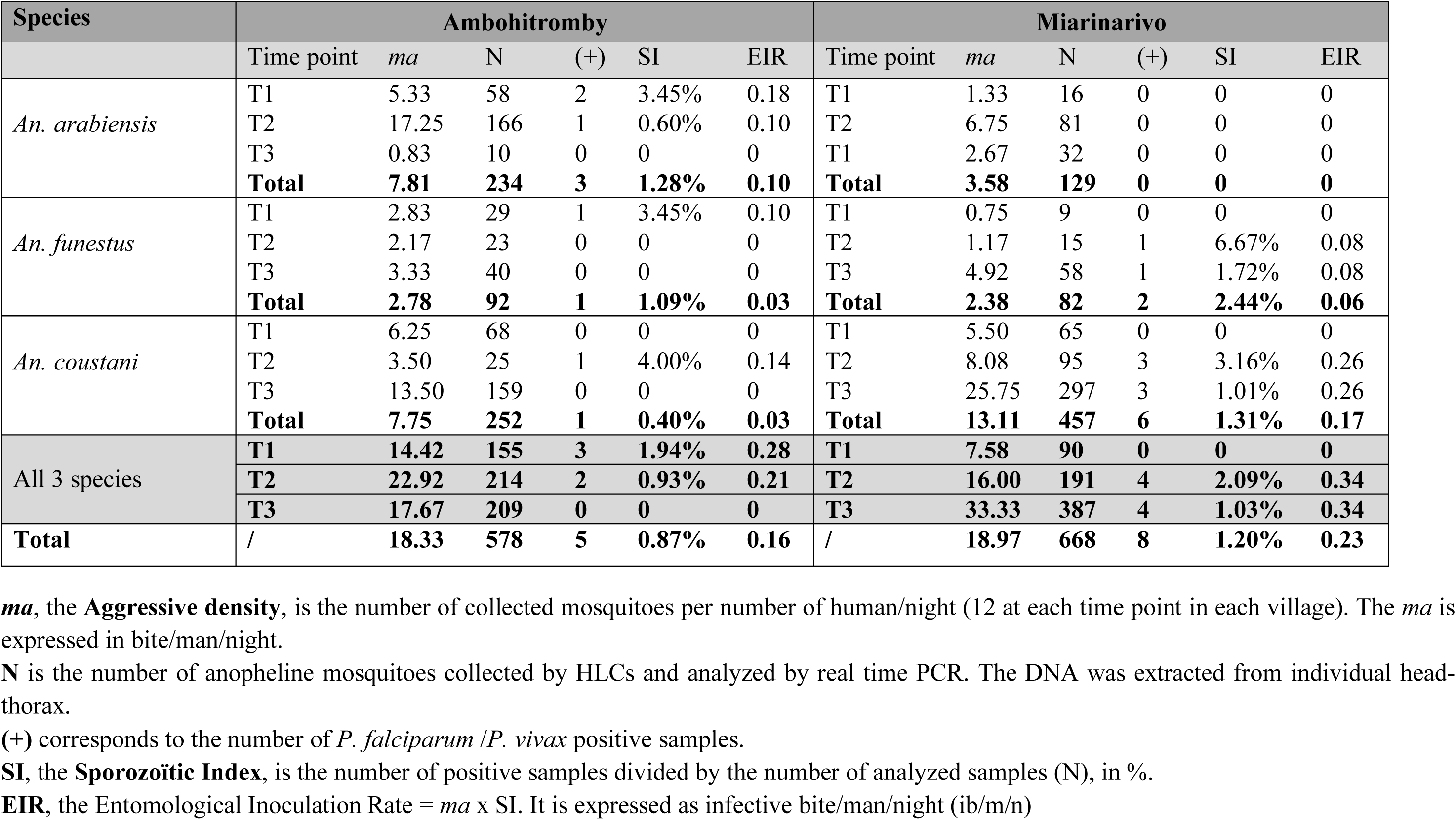
Entomological parameters of the malaria vectors confirmed positive by the SYBR Green assay, at the three time points.

**Fig 5.**
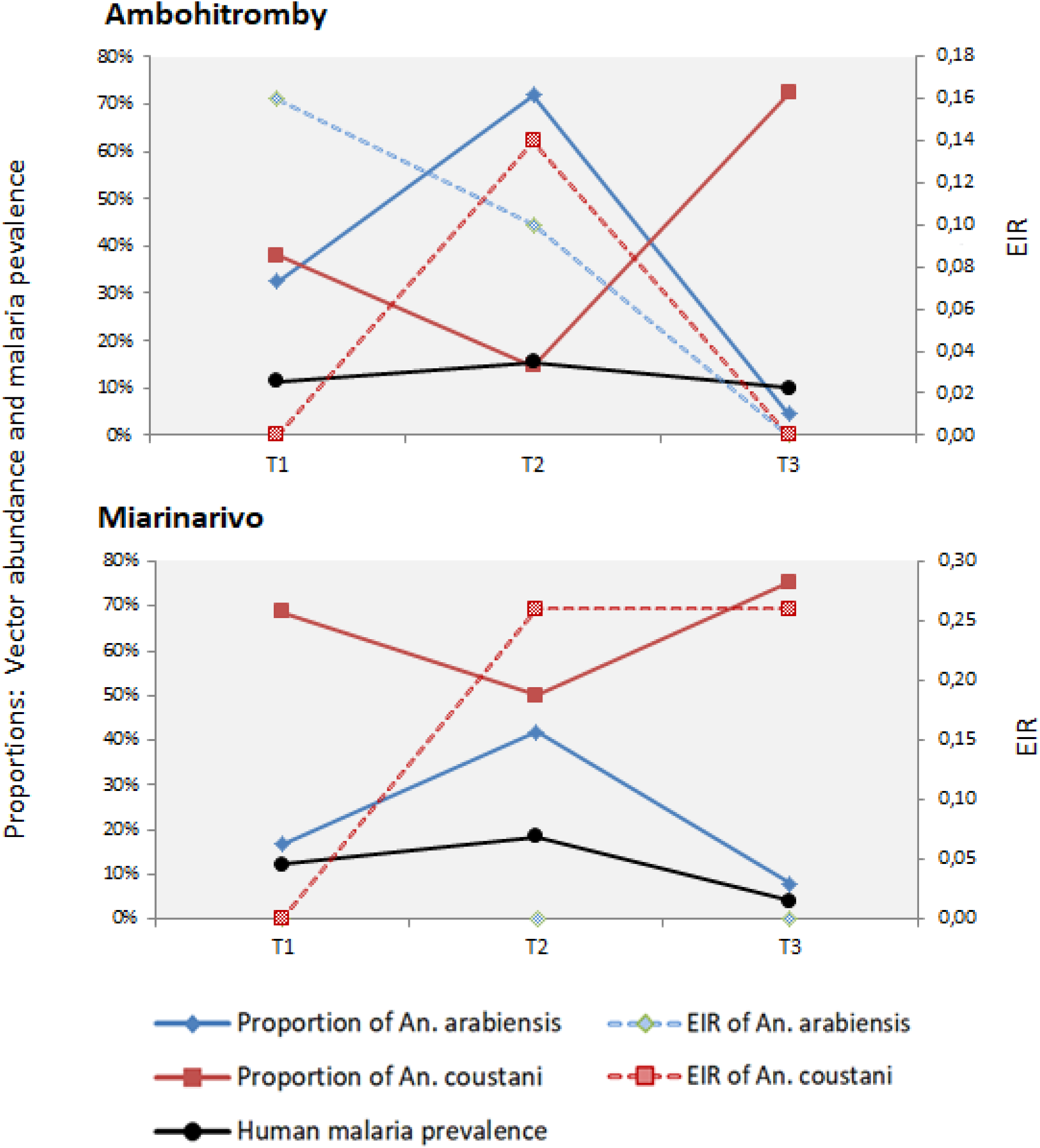
Variation of the relative abundance and EIR of *An. arabiensis* and *An. coustani* in Ambohitomby and Miarinarivo.

In conclusion, our data show that in close by villages with similar malaria prevalence in the human population malaria transmission involved two different mosquito species and notably involved *An. coustani* as the major vector in Miarinarivo. Detailed analysis of the EIR over time (Table 6) shows that malaria transmission occurred at the beginning and middle malaria transmission in Ambohitromby due to *An. arabiensis*, while occurring at the mid-course and vanishing of the malaria transmission season in Miarinarivo, due to *An. coustani*. Overall, the population in Ambohitromby are expected to receive 24 infected bites per person over the malaria season (December – April) and 34.5 infected bites per person in Miarinarivo.

The *Plasmodium* carriage rate in mosquitoes was not significantly different between the two villages (Pearson’s test *P*=0.546, Pearson Chi2 Test).

## Discussion

The objective of the study was to estimate the level of malaria transmission and identifying the mosquito species involved in two close by villages in a region of Madagascar where malaria is still a high public health problem. Indeed, in that region no such study had ever been conducted despite the high number of malaria patients attending the local health center. To our knowledge, the only study carried out at the same site just provided entomological data and goes back to 1992 [34].

### Similarity in human malaria prevalence in the two villages

Parasitological data in the asymptomatic villagers revealed malaria infection cases due mainly to *P. falciparum*, in addition to *P. vivax* and *P. malariae*. The global malaria prevalence over the whole study was 11.9%, as determined by PCR, and 4.8% by microscopy. These values are similar to the ones reported in the Tsiroanomandidy study performed in March 2014 [13]. Our results highlight the high prevalence of submicroscopic *Plasmodium* carriage as well which represents 50 to 75 % of the investigated cases (negative by microscopy, positive by PCR). Our data show similar malaria prevalence between the two villages. This is in sharp contrast to the significant difference that was observed in 2016 between the school aged children of Ambohitromby and of Miarinarivo (unpublished data). Indeed, in 2016 a significant difference (*P*=0.019, Chi2 test) in malaria RDT prevalence was observed between Ambohitromby (19.5%, n=41) and Miarinarivo (6.3%, n=96), but not in 2017 (this study) nor 2018 (Bourgouin and coll. unpublished, Fig 6). This variation in malaria prevalence across years in the two villages might result from better mosquito net coverage of the populations or climatic and ecological changes impacting *Anopheles* density [35,36]. However, no mosquito net distribution was done between 2016 and 2018 in Andriba. Therefore, it might be possible that in Ambohitromby 2016, the local conditions facilitated the development of an increased number of mosquito breeding sites leading to increased *Anopheles* vector population size and subsequent increased transmission.

**Fig 6.**
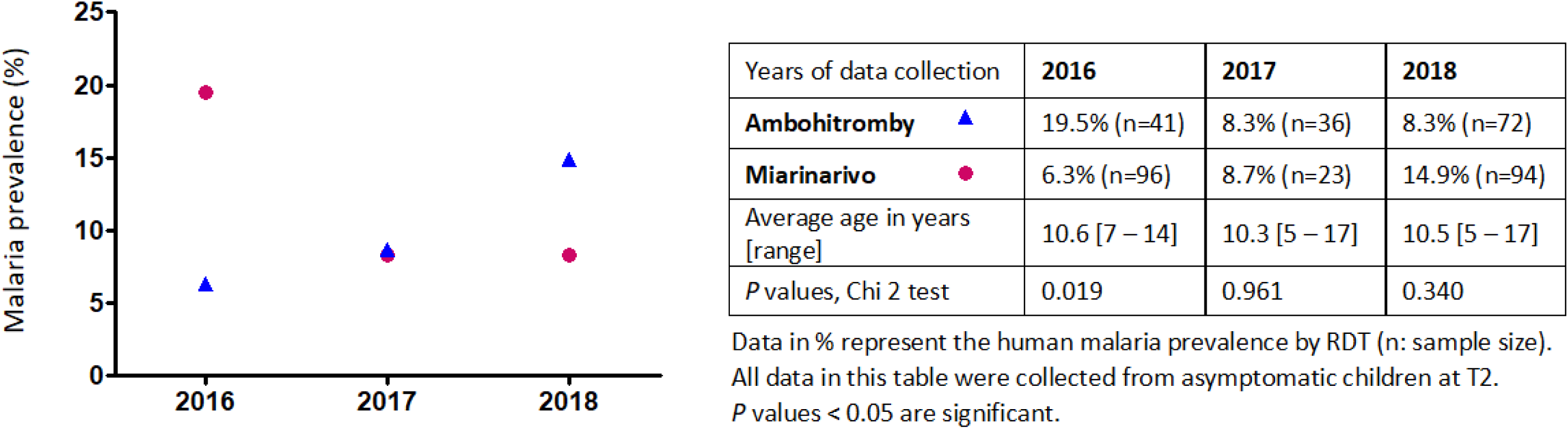
School children malaria prevalence in Ambohitromby and Miarinarivo between 2016 and 2018. Data were obtained from RDT testing of school age children in March each year.

### Vector species diversity and behavior in the two villages

Entomological data from HLCs and PSCs, showed that the overall abundance and mosquito species diversity were similar between the two villages surveyed. Among the known malaria vectors, only *An. funestus* showed a strong tendency for endophily for both biting and resting, not departing from its known behavior in Andriba [34] and in other African regions including Madagascar [5,18,27,28,30,37]. Across the malaria transmission season, from December to April, we recorded differences in the aggressive density according to the time point. *An. arabiensis* was the most aggressive species at the mid-term of the transmission season (February) in Ambohitromby while it was *An. coustani* the most aggressive species at the onset (December) and at the late-term (April) in each village. This change in species aggressiveness over the malaria transmission season could be explained by changes in the ecological environment, precisely the rice fields. Indeed, the growing phases of the rice determine important changes in the characteristics of the *Anopheles* breeding sites [38]. Fields with rice in the early stages of growth and with young short plants offer sunny breeding sites favorable for the development of *An. gambiae* s.l. larvae. As the rice plants grow, they shade the water of the rice fields which become less favorable for *An. gambiae* s.l. larvae, giving way to *An. coustani* larvae that prefer shaded breeding sites. This transition from *An. gambiae* s.l. to *An. coustani* in breeding sites by increasing vegetation cover was also demonstrated for borrow pits in Ethiopia [39]. Through the analysis of the *Anopheles* vector feeding behavior, it is noticeable that both *An. arabiensis* and *An. coustani* exhibited a significant different feeding behavior in each village towards increased endophagy in Miarinarivo (Table 4). This observation might be the signature of different populations within each species. It cannot be excluded though that the local environment as distance between houses and breeding sites, their number and size, and the structure of the villages itself contribute to the observed differences in *An. arabiensis* and *An. coustani* endophagy between the two villages.

### Different contribution of *Anopheles* species to malaria transmission between the two villages

*P. falciparum* and *P. vivax* sporozoites were detected by PCR in only three anopheline species: *An. funestus, An. arabiensis* and *An. coustani*. All mosquitoes positive for *Plasmodium* were collected by HLCs, with 8/13 collected outdoor, highlighting that malaria transmission in Andriba was mainly occurring outdoor. Whereas *An. funestus* and *An. arabiensis* are well known malaria vector in Madagascar, the contribution of *An. coustani* to malaria transmission has been suspected on several occasions due to its high density and propency to anthropophily [31]. It was only recently that some *An. coustani* samples were detected CSP-positive by ELISA [9]. Our data using real time PCR, clearly demonstrated the vector role of *An. coustani* in malaria transmission in Andriba. *An. coustani* is also known to be a malaria vector in continental Africa: in Cameroon [40], Zambia [41], Kenya [42], Ethiopia [39].

Surprisingly, our data revealed that *An. coustani*, was the major malaria vector in Miarinarivo, despite the presence of *An. funestus* and *An. arabiensis*. In that village, people were exposed to 25.5 infected bites from *An. coustani* across the time period survey, compared to only 4.5 infected bites in Ambohitromby, despite its relatively high abundance in this latter village. Infected *An. coustani* were captured equally outdoor and indoor in Miarinarivo. By contrast, *An. arabiensis* was the major vector in Ambohitromby, responsible for 15 infected bites across the same time period while not involved at all in malaria transmission in Miarinarivo. *An. arabiensis* infected mosquitoes were found both in indoor and outdoor HLCs. *An. funestus* contributed to a minor extent to malaria transmission in both villages being responsible for 4.5 and 9 infected bites during the whole survey in Ambohitromby and Miarinarivo respectively. Infected *An. funestus* were captured both indoor and outdoor.

The different contribution of malaria vector species might result from a different layout of the houses in each village and their distance from the rice fields as previously argued [43]. Indeed, the satellite view of the two villages shows that the houses in Miarivaniro are more numerous and very close to each other compare to houses in Ambohitromby, and that Miarinarivo is surrounded by more and closer rice fields, which is particularly favorable to the large number of *An. coustani* recorded in Miarinarivo.

Our work revealed also that two out of 40 *An. rufipes* analyzed by TaqMan assay were found possibly carrying *Plasmodium* sporozoites, although we were not able to identify the *Plasmodium* species using the SYBR Green assay which is known to be less sensitive than the TaqMan assay. To date, *An. rufipes* has never been reported naturally infected with *Plasmodium* in Madagascar. In continental Africa, it was found naturally infected with *P. falciparum* in Burkina Faso [31,44] and more recently in Cameroon [45]. Lastly, none of the *An. mascarensis* samples (n=59) were found positive for *Plasmodium* although it is known as a malaria vector in other Malagasy areas [8,10,46,47].

Overall, this study demonstrates the variability of vector biology dynamics between two neighboring villages with similar ecological setting. Our data can be used to better describe the epidemiology and transmission of malaria in Madagascar and to provide relevant information to guide for adapted malaria control. In an epidemiological context such as Madagascar, marked by the prevalence of *P. falciparum* and *P. vivax* and the presence of several vector species, understanding vector-specific contributions to the transmission of these two main *Plasmodium* species constitutes a malaria elimination challenge.

## Acknowledgements

We thank lab technicians from the Institut Pasteur de Madagascar who helped in data collection and experimentations: Mandaniaina R. Andriamiarimanana, Emma Rakotomalala, Rado L. Rakotoarison, Rakotoniaina M. Tanjona and Maminirina F. Ambinintsoa. We also thank Richard Paul from the Institut Pasteur who helped in statistical analyzes and Miriam K. Laufer from the University of Maryland School of Medicine for critical reading of the manuscript.

## Additional information

### Related manuscripts

The authors do not have a related or duplicate manuscript under consideration (or accepted) for publication elsewhere.

### Ethics approval and consent to participate

This study followed ethical principles according to the Helsinki Declaration and was approved by the Malagasy Ethical Committee of the Ministry of Health (agreements N°122-MSANP/CE– 2015 and N°141-MSANP/CE–2014).

### Availability of data and materials

Data generate or analyzed during this study are included in this published article. Datasets used during the current study are available from the corresponding author on reasonable request.

## List of abbreviations

ACT: artemisinin-based combination therapy;
CI: confidence interval;
CSP: circumsporozoite protein;
DNA: deoxyribose nucleic acid;
EIR: entomological inoculation rate;
HLCs: human landing catches;
HN: human-night;
ib/n/m: infective bite per man per night;
IRS: insecticide residual spraying;
PBS: phosphate-buffered saline;
PCR: polymerase chain reaction;
PSCs: pyrethrum spray catches;
RDT: rapid diagnostic test;
RNA: ribonucleic acid;
SI: sporozoite index.

