## Supplementary information for "Differential contribution of *Anopheles coustani* and *Anopheles arabiensis* to the transmission of *Plasmodium falciparum* and *Plasmodium vivax* in two neighboring villages of Madagascar"

### Supporting information

**S1 Table.** **Sequences of the primers and TaqMan probes used for the morphological identification of *An. gambiae*/*An. arabiensis* and for *Plasmodium* detection in *Anopheles* mosquitoes.**

| **Species** | **Primer/Probe** | **Sequence** | **Reference** |
| --- | --- | --- | --- |
| *An. gambiae* *s.l.* | UNI-F | 5'-GTGAAGCTTGGTGCGTGCT-3' | Walker *et al.,* 2007 |
| *An. gambiae* *s.l.* | UNI-R | 5'-GCACGCCGACAAGCTCA-3' |  |
| *An. gambiae* | Probe | 5'-VIC- CGGTATGGAGCGGGACACGTA-MGB |  |
| *An. arabiensis* | Probe | 5'-6FAM-TAGGATGGAGAAGGACACTTA-MGB |  |
| *Plasmodium* *spp.* | Plasmo1-F | 5'-GTT AAG GGA GTG AAG ACG ATC AGA | Modified from Rougemont *et al*., 2004 |
|  | Plasmo2-R | 5'-AAC CCA AAG ACT TTG ATT TCT CAT AA |  |
|  | Plasmoprobe | 5'-NED-TCGTAATCTTAACCATAAAC-MGB |  |
| *P. falciparum*- Cytb | Cytb F | 5’-ATGGATATCTGGATTGATTTTATTTATGA | Canier *et al.*, 2013 |
|  | Cytb R | 5’- TCCTCCACATATCCAAATTACTGC |  |
| *P. vivax*-Cytb | Cytb F | 5’- TGCTACAGGTGCATCTCTTGTATTC |  |
|  | Cytb R | 5’- ATTTGTCCCCAAGGTAAAACG |  |

**S2 Table. Population by age group and Sex**

|  | | **Ambohitromby** | **Miarinarivo** | **Total** |
| --- | --- | --- | --- | --- |
| Sample size | | 218 | 162 | 380 |
| Age group | <5 | 34 | 28 | 62 (16.32%) |
|  | [5-10[ | 43 | 24 | 67 (17.63%) |
|  | [10-15[ | 46 | 23 | 69 (18.16%) |
|  | >15 | 95 | 87 | 182 (47.89%) |
| Sex | Male | 97 | 75 | 172 (45.26%) |
|  | Female | 121 | 87 | 208 (54.73%) |

The numbers in parenthesis are the proportions.

**S3 Table. Mosquitoes collected by HLCs inside and outside houses, in Ambohitromby and Miarinarivo over the 3 survey time points.**

| **Mosquito species** | **Ambohitromby** | | | | | | | | **Miarinarivo** | | | | | | | |
| --- | --- | --- | --- | --- | --- | --- | --- | --- | --- | --- | --- | --- | --- | --- | --- | --- |
| Time points | **T1** | | **T2** | | **T3** | | **Total** | | **T1** | | **T2** | | **T3** | | **Total** | |
| Place of collection (Ind/Out) | Ind | Out | Ind | Out | Ind | Out | **Ind** | **Out** | Ind | Out | Ind | Out | Ind | Out | **Ind** | **Out** |
| *Anopheles arabiensis* | 19 | 45 | 49 | 158 | 3 | 7 | **71** | **210** | 9 | 7 | 30 | 51 | 16 | 16 | **55** | **74** |
| *Anopheles gambiae* | 0 | 0 | 1 | 3 | 0 | 0 | **1** | **3** | 1 | 0 | 2 | 0 | 1 | 0 | **4** | **0** |
| *Anopheles funestus* | 14 | 20 | 13 | 13 | 31 | 9 | **58** | **42** | 8 | 1 | 7 | 7 | 29 | 30 | **44** | **38** |
| *Anopheles mascarensis* | 4 | 20 | 1 | 12 | 0 | 12 | **5** | **44** | 1 | 4 | 0 | 2 | 1 | 9 | **2** | **15** |
| *Anopheles coustani* | 12 | 63 | 1 | 41 | 3 | 159 | **16** | **263** | 21 | 45 | 37 | 60 | 80 | 229 | **138** | **334** |
| *Anopheles squamosus/cydippis* | 1 | 12 | 0 | 43 | 3 | 26 | **4** | **81** | 0 | 8 | 19 | 21 | 4 | 11 | **23** | **40** |
| *Anopheles rufipes* | 2 | 7 | 2 | 8 | 0 | 0 | **4** | **15** | 2 | 3 | 6 | 5 | 4 | 5 | **12** | **13** |
| *Anopheles maculipalpis* | 0 | 7 | 0 | 9 | 0 | 0 | **0** | **16** | 5 | 5 | 1 | 8 | 2 | 3 | **8** | **16** |
| *Anopheles pretoriensis* | 0 | 0 | 0 | 0 | 0 | 0 | **0** | **0** | 0 | 0 | 0 | 0 | 1 | 0 | **1** | **0** |
| *Culex antennatus* | 3 | 27 | 38 | 220 | 0 | 4 | **41** | **251** | 26 | 23 | 35 | 64 | 4 | 21 | **65** | **108** |
| *Culex quinquefasciatus* | 1 | 4 | 40 | 63 | 0 | 0 | **41** | **67** | 4 | 11 | 8 | 15 | 1 | 18 | **13** | **44** |
| *Culex bitaeniorhyncus* | 0 | 0 | 0 | 0 | 0 | 0 | **0** | **0** | 0 | 1 | 0 | 1 | 0 | 0 | **0** | **2** |
| *Culex univittatus* | 0 | 4 | 0 | 0 | 0 | 0 | **0** | **4** | 0 | 0 | 0 | 0 | 0 | 0 | **0** | **0** |
| *Culex decens* | 0 | 1 | 0 | 0 | 0 | 0 | **0** | **1** | 1 | 1 | 0 | 0 | 0 | 0 | **1** | **1** |
| *Culex giganteus* | 0 | 0 | 0 | 0 | 0 | 0 | **0** | **0** | 1 | 1 | 2 | 2 | 0 | 2 | **3** | **5** |
| *Mansonia uniformis* | 1 | 7 | 2 | 7 | 1 | 2 | **4** | **16** | 2 | 3 | 4 | 21 | 4 | 30 | **10** | **54** |
| *Aedes skusea* | 0 | 5 | 0 | 0 | 0 | 0 | **0** | **5** | 1 | 2 | 0 | 0 | 0 | 0 | **1** | **2** |
| *Aedes vittatus* | 0 | 1 | 0 | 1 | 0 | 0 | **0** | **2** | 0 | 0 | 0 | 0 | 0 | 0 | **0** | **0** |
| *Aedes tiptoni* | 0 | 1 | 0 | 0 | 0 | 0 | **0** | **1** | 2 | 6 | 0 | 0 | 1 | 0 | **3** | **6** |
| *Aedes albopictus* | 0 | 0 | 0 | 1 | 0 | 0 | **0** | **1** | 1 | 0 | 1 | 0 | 0 | 0 | **2** | **0** |
| *Aedes circumlateolus* | 0 | 0 | 0 | 0 | 0 | 0 | **0** | **0** | 0 | 0 | 0 | 0 | 0 | 1 | **0** | **1** |
| *Coquillettidia grandidieri* | 0 | 0 | 0 | 0 | 0 | 0 | **0** | **0** | 0 | 0 | 0 | 0 | 0 | 2 | **0** | **2** |
| **Grand total** | 57 | 224 | 146 | 576 | 41 | 219 | **245** | **1022** | 84 | 121 | 150 | 257 | 147 | 377 | **385** | **755** |

Ind: Indoor; Out: Outdoor

**S4 Table. Proportion of mosquitoes collected resting indoor by PSC (Endophilic rate).**

| **Species** | **Ambohitromby** | | | | **Miarinarivo** | | | | **Grand total** |
| --- | --- | --- | --- | --- | --- | --- | --- | --- | --- |
| Time points | T1 | T2 | T3**^a^** | **Total** | T1 | T2**^b^** | T3 | **Total** |  |
| *An. arabiensis* | 9 | 3 | 0 | **12**  (31.57%) | 0 | 0 | 0 | **0** | **12**  (18.75%) |
| *An. funestus* | 10 | 4 | 2 | **16**  (42.10%) | 5 | 2 | 18 | **25**  (96.15%) | **41**  (64.06%) |
| *An. mascarensis* | 3 | 0 | 1 | **4**  (10.52%) | 1 | 0 | 0 | **1**  (3.84%) | **5**  (7.81%) |
| *An. coustani* | 2 | 0 | 4 | **6**  (15.78%) | 0 | 0 | 0 | **0** | **6**  (9.37%) |
| *An. rufipes* | 1 | 0 | 0 | **1** | 0 | 0 | 0 | 0 | **1** |
| *Cx antennatus* | 0 | 0 | 0 | **0** | 0 | 2 | 0 | **2** | **2** |
| *Cx quinquefasciatus* | 0 | 3 | 0 | **3** | 0 | 0 | 0 | **0** | **3** |
| **Total** | **25** | **10** | **7** | **42** | **6** | **4** | **18** | **28** | **70** |

**^a^** Captures were performed during 2 days on 3. **^b^** Only 2 houses on 5 were performed. Values in brackets represent the relative abundance (endophilic rate) to known malaria vectors collected resting indoor (*An. arabiensis, An. funestus, An. mascarensis* and *An. coustani*), for a total of 38 mosquitoes in Ambohitromby and 26 in Miarinarivo. 3 PSCs were performed at each time point per village, except in Ambohitromby at T3 where only 2 could be performed; this led to a total of 17 PSCs, 8 in Ambohitromby and 9 in Miarinarivo.
